# Vpr Co-assembles with Gag during HIV-1 Assembly

**DOI:** 10.1101/2020.03.23.004689

**Authors:** Kate Bredbenner, Sanford M Simon

## Abstract

The HIV-1 accessory protein Vpr is packaged into new virions at a 7:1 ratio of Gag/Vpr. Previous biochemical and genetic analysis has shown that Vpr gets packaged into virions via an LXXLF motif on the p6 domain of the Gag structural polyprotein. The kinetics of Vpr packaging compared to Gag assembly was previously unknown. Here, we confirm via biochemistry and imaging that fluorescently tagged Vpr gets packaged into virus-like particles only when the LXXLF motif is intact. When the LXXLF motif is mutated, Vpr is no longer recruited to Gag assemblies. When Vpr and Gag assembly are imaged together, we see that Vpr co-assembles with a slight delay compared to Gag suggesting that Vpr is not being recruited to the membrane with Gag but is instead being recruited to actively assembling Gag.

**Importance:** HIV-1 affects over 30 million people around the globe, and although we have good treatments, there is still no cure. The virus encodes 15 distinct proteins, and four of those proteins are known as accessory proteins. Vpr is one of the accessory proteins that is packaged into HIV-1 by interacting with the Gag structural protein. Without Vpr, HIV-1 is not as infectious. Our research shows that Vpr is packaged into new viruses as the virus is being formed rather than being put in towards the end of the assembly of a virus. Getting a clearer view of each step in the process of assembling each virion will help inform future treatments and help with overall comparisons between the assembly of different viruses.

## Introduction

HIV-1 viruses assemble on the plasma membrane of infected cells via polymerization of the p55Gag structural polyprotein (1). It takes between 5-25 minutes between the arrival of the HIV-1 genome and assembly of the Gag protein (2,3). Each virus contains approximately 2400 Gag polyproteins (4). After recruitment of Gag is complete, host-cell ESCRT proteins are recruited transiently to the sites of assembly in order to help the particle scission from the cell (5–8). Within the p6 domain of Gag, the PTAP motif recruits TSG101(9,10) and the YPXL motif recruits Alix (11–14) which are both necessary for viral budding and release (9,12,15). Alix also helps recruit other downstream ESCRT components necessary for scission such as CHMP4B and VPS4A (8,11,13,14).

HIV-1 also encodes several other proteins that are packaged into virions. The Pol proteins (protease, reverse transcriptase, and integrase) are synthesized and packaged via a fusion to Gag. They are made as a consequence of a −1 ribosomal frameshift that occurs approximately 5% of the time resulting in approximately 120 copies of the Pol proteins per virion(16). HIV-1 also encodes several additional accessory proteins such as Vif, Nef, and Vpr that are packaged into virions.

Vpr is a 97-amino acid accessory protein that is packaged into virions via interaction with the LXXLF motif located in the p6 domain of Gag (17,18). The LXXLF motif on p6 is further towards the carboxy terminus than both the PTAP and the YPXL motifs which recruit the ESCRT machinery. Vpr is packaged into each virus with a ratio of Gag/Vpr of 7:1, which corresponds to approximately 340 molecules per virion (19–22). Although Vpr is not necessary for HIV-1 growth in culture, it might play a role during *in vivo* replication (23,24). Proposed functions of Vpr include influences on nuclear entry of viral proteins (25,26) and induction of G2/M cell cycle arrest or cell apoptosis (27–29).

Because of its ability to interact with Gag and be packaged in-trans, Vpr has been a popular target to use for fusions to deliver proteins into viruses. Previous research has used Vpr to deliver the HIV-1 protease, integrase, and reverse transcriptase proteins into virions outside of their typical packing as fusions to Gag (30–34). Other research has packaged the TEV protease or GFP into virions with Vpr (35–39).

The kinetics of the recruitment of Vpr and packaging into viruses is an open question. To determine the kinetics, we fused an mEGFP to Vpr and confirmed that it was packaged into virus-like particles (VLPs). We also confirmed that a premature truncation of the Gag polyprotein, resulting in a partial truncation of the LXXLF motif, prevented packaging of Vpr, but still allowed assembly and release of VLPs. The recruitment of Vpr during assembly of VLPs shows that it recruits with a slight delay compared to Gag.

## Results

### Vpr is incorporated into VLPs

To determine the kinetics of Vpr recruitment, we wanted to be able to image Vpr using fluorescence. Thus, we first had to confirm that fluorescently tagged Vpr was packaged specifically into our VLPs. We created an mEGFP-Vpr construct and co-expressed it with an NL4.3-GagPol packaging vector with an mCherry inserted after the matrix domain of Gag (GagPol-mCherry) (Fig 1). This latter construct will make mostly Gag-mCherry, but also GagPol-mCherry when slippage occurs. The expression and packing of the mEGFP-Vpr into VLPs was confirmed by Western blots (Fig 2A). To confirm that Vpr was specifically being recruited to sites of assembly via the LXXLF motif, we introduced a stop codon into the LXXLF motif of Gag which removed the last 9AA of p6 while leaving the Alix and TSG101 recruitment motifs intact (GagPol-mCherry-LXXLF) (Fig 1). This mutation also leaves the Pol amino acid sequence fully intact. When we co-expressed mEGFP-Vpr with GagPol-mCherry-LXXLF, we were able to collect VLPs that that were positive for capsid and mCherry, albeit at reduced levels, but mEGFP-Vpr was not detectable (Fig 2A).

**Figure 1.**
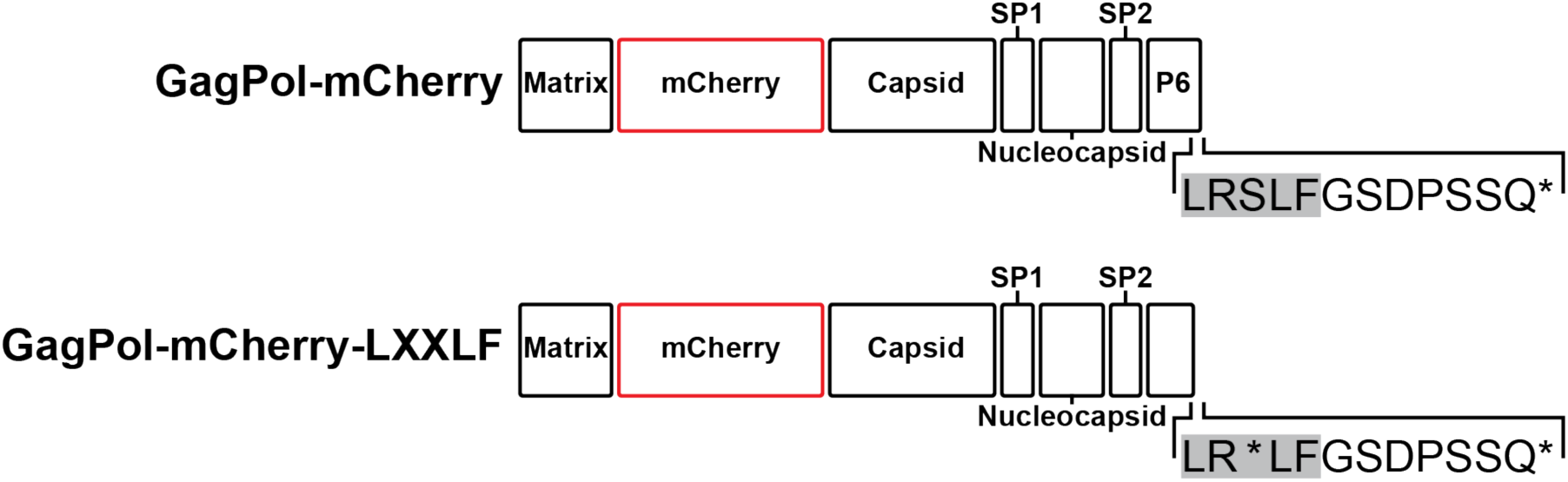
Schematic of GagPol Plasmids. The sequences for GagPol-mCherry and GagPol-mCherry-LXXLF were constructed from NL4.3 sequences which have the slippage site intact. The mCherry is inserted in each construct, without a linker, at the end of matrix domain prior to the Matrix/Capsid cleavage site. GagPol-mCherry-LXXLF has a stop codon introduced in the middle of the LXXLF Vpr recruitment motif which removes the last 9 amino acids from Gag but leaves the Pol coding sequence intact.

**Figure 2.**
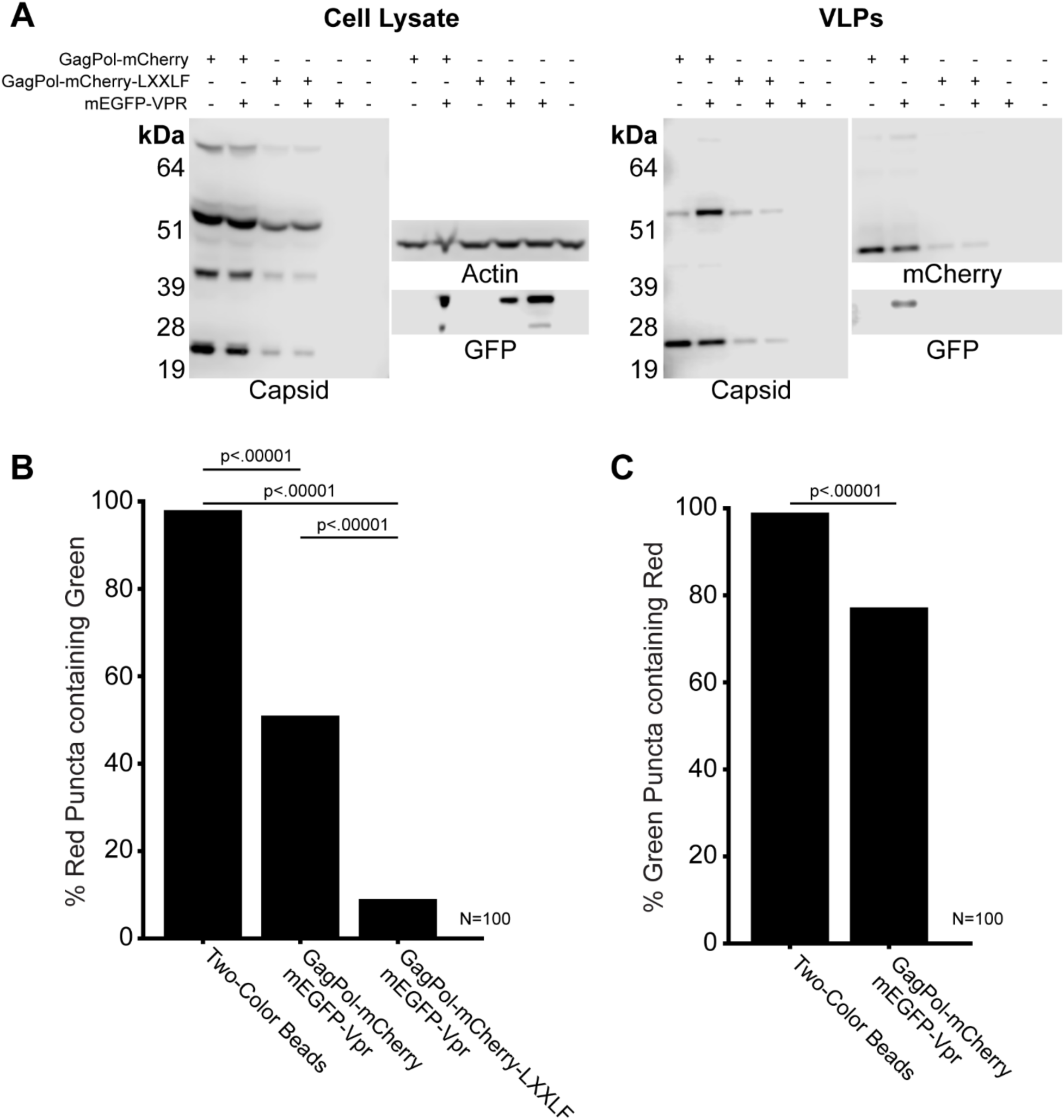
Vpr is packaged specifically into VLPs. **A** Cell lysate (left) and VLPs (right) collected from Hek293T cells 48 hours after transfection. Cell lysate was probed with anti-capsid, anti-actin, and anti-GFP antibodies. Western blots were probed with anti-capsid, anti-mCherry, and anti-GFP antibodies. **B** shows the quantification of the percent of red puncta that were also green (n=100). **C** shows the quantification of the percent of green puncta that were also red (n=100).

To further test the assembly of mEGFP-Vpr into VLPs, we co-transfected it together with GagPol-mCherry and collected supernatant after 24 hours. There are some potential problems with this approach. First, VLPs are created via transient transfection, which means that the VLPs come from a population of cells with different amounts of plasmid and expressing differing levels of each protein. Further, the collection of VLPs includes small pieces of cell debris that go through our filter. For this reason, quantifying the relative levels of proteins VLPs collected from a mixed population of cells has potential limitations for quantifying the packaging of Vpr. Additionally, VLPs with GagPol-mCherry can be excited at 488 nm. While they will not be as efficiency excited as a GFP, at a high concentration VLPs with GagPol-mCherry excited at 488nm may falsely be recorded as positives for mEGFP-Vpr. To limit any false positives, we imaged VLPs (n=100) with GagPol-mCherry and GagPol-mCherry-LXXLF in the absence mEGFP-Vpr to quantify the cross-channel contamination so the mCherry excited by the 488nm laser can be subtracted from the mEGFP images.

Despite the stringent subtraction of the mCherry signal from the GFP channel and the inherent problems with collecting VLPs from transient transfection, 51% of all GagPol-mCherry VLPs also contained detectable mEGFP-Vpr (Fig 2B). When we analyzed VLPs made with GagPol-mCherry-LXXLF which is missing the Vpr recruitment domain, the percent of GagPol-mCherry VLPs which contained detectable mEGFP-Vpr dropped to 9% (Fig 2B).

To test if mEGFP-Vpr was only found in VLPs and not found in either other cell debris or in puncta without Gag, we checked mEGFP-Vpr positive puncta (n=100) and found that 77% of them were also GagPol-mCherry positive (Fig 2C). When we transfect cells with GagPol encoding an embedded fluorescent protein, we co-transfect with a plasmid with GagPol without a tag. This avoids morphological problems in VLPs that occur when all of the copies of GagPol are expressed as fusion proteins (40). The ratio we use is a 1:4 ratio of tagged/untagged GagPol. We believe that the percentage of mEGFP-Vpr puncta that also contain GagPol-mCherry is not 100% due to a proportion of cells that were only transfected with untagged GagPol and mEGFP-Vpr without tagged GagPol. The results from Western blots and imaging are consistent with mEGFP-Vpr being packaged specifically into VLPs. We also confirm that the LXXLF motif is necessary for that recruitment.

### Vpr co-assembles with GagPol

We next examined the recruitment of mEGFP-Vpr in cells expressing either GagPol-mCherry or GagPol-mCherry-LXXLF. Still images from cells with GagPol-mCherry and mEGFP-Vpr six hours after transfection showed puncta in both red and green that co-aligned. In contrast, when we co-transfected the GagPol-mCherry-LXXLF with the mEGFP-Vpr, cells assembled VLPs, but mEGFP-Vpr was not detected at sites of GagPol (Fig 3A). Quantification of VLPs in the still images showed that 78% of the selected GagPol-mCherry puncta on the cell were also mEGFP-Vpr positive, but only 5% of GagPol-mCherry-LXXLF puncta were mEGFP-VPR positive (Fig 3B). The percentage of GagPol-mCherry puncta that were positive for mEGFP-Vpr on the surface of cells was higher than from isolated VLPs. This substantiates the conclusion that a contributing factor to the lower level of colocalization in the VLPs were cells expressing GagPol-mCherry but not mEGFP-Vpr.

**Figure 3.**
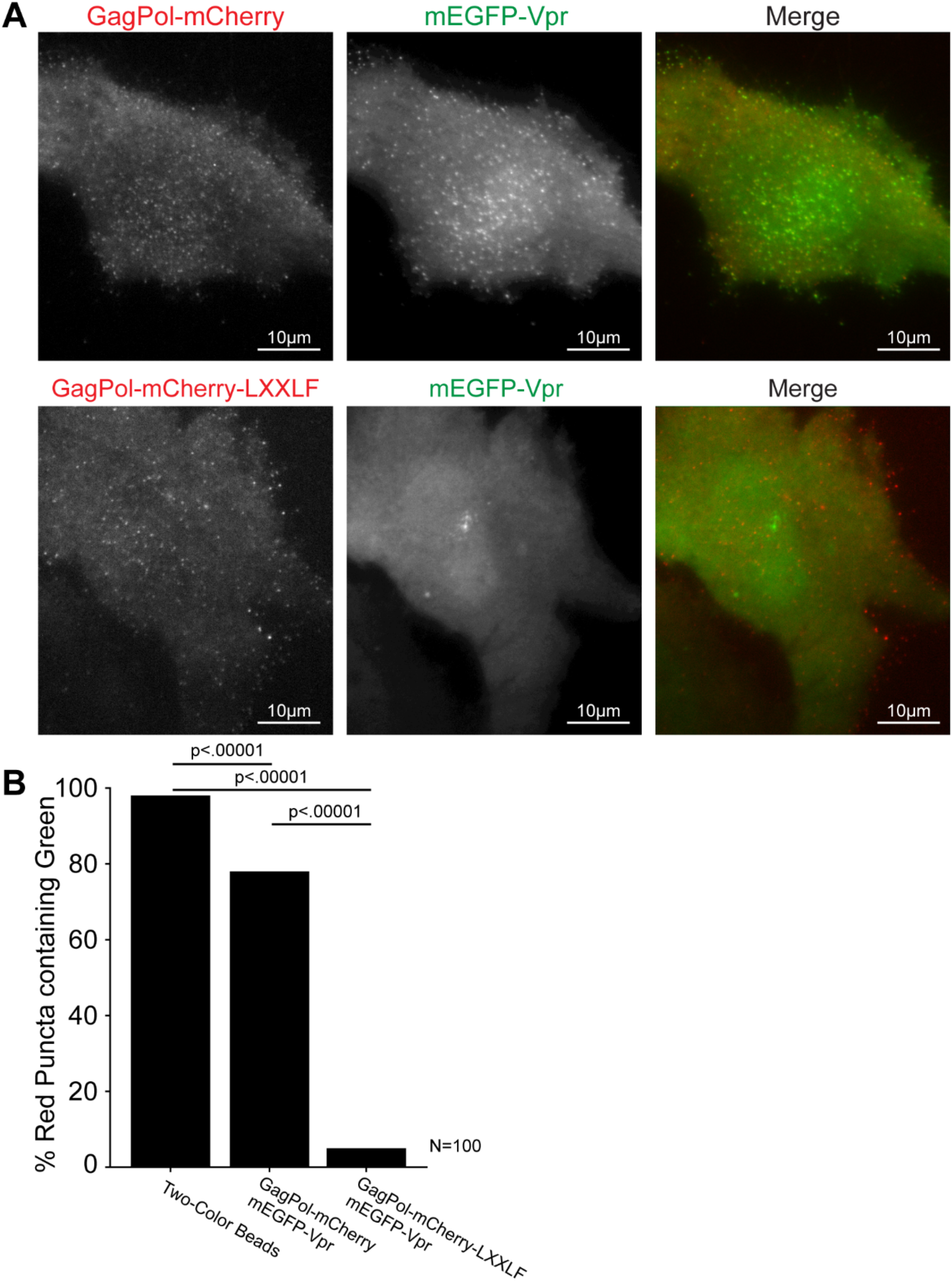
GagPol with or without the LXXLF mutation. Single frames from a sequence of HeLa cells actively assembling new particles (**A**). The top row of images is from cells transiently expressing GagPol-mCherry and mEGFP-Vpr. The bottom row is from cells transiently expressing GagPol-mCherry-LXXLF and mEGFP-Vpr. The first column is GagPol in red, the second is Vpr in green, and the final column is a merge of the first two. All scale bars are 10µm. Quantification of puncta from two-color beads and GagPol-mCherry or GagPol-mCherry-LXXLF puncta on from cells (n=100).

To determine the kinetics of recruitment of Vpr to VLPs, we imaged mEGFP-Vpr in live cells. When we did a time lapse imaging of mEGFP-Vpr together with GagPol-mCherry in HeLa cells, we saw that mEGFP-Vpr co-localized with GagPol (Fig 4A). The intensity of the fluorescence from mEGFP-Vpr reaches a plateau, indicating the end of net recruitment, similar to the plateau reached by GagPol assembly. Vpr does show a delay of 4-6 minutes in reaching its plateau when compared to GagPol. The kinetics or recruitment we observed of the mEGFP-Vpr was the same whether we used only untagged GagPol or a mixture with untagged and tagged GagPol-mCherry (Fig 4B).

**Figure 4.**
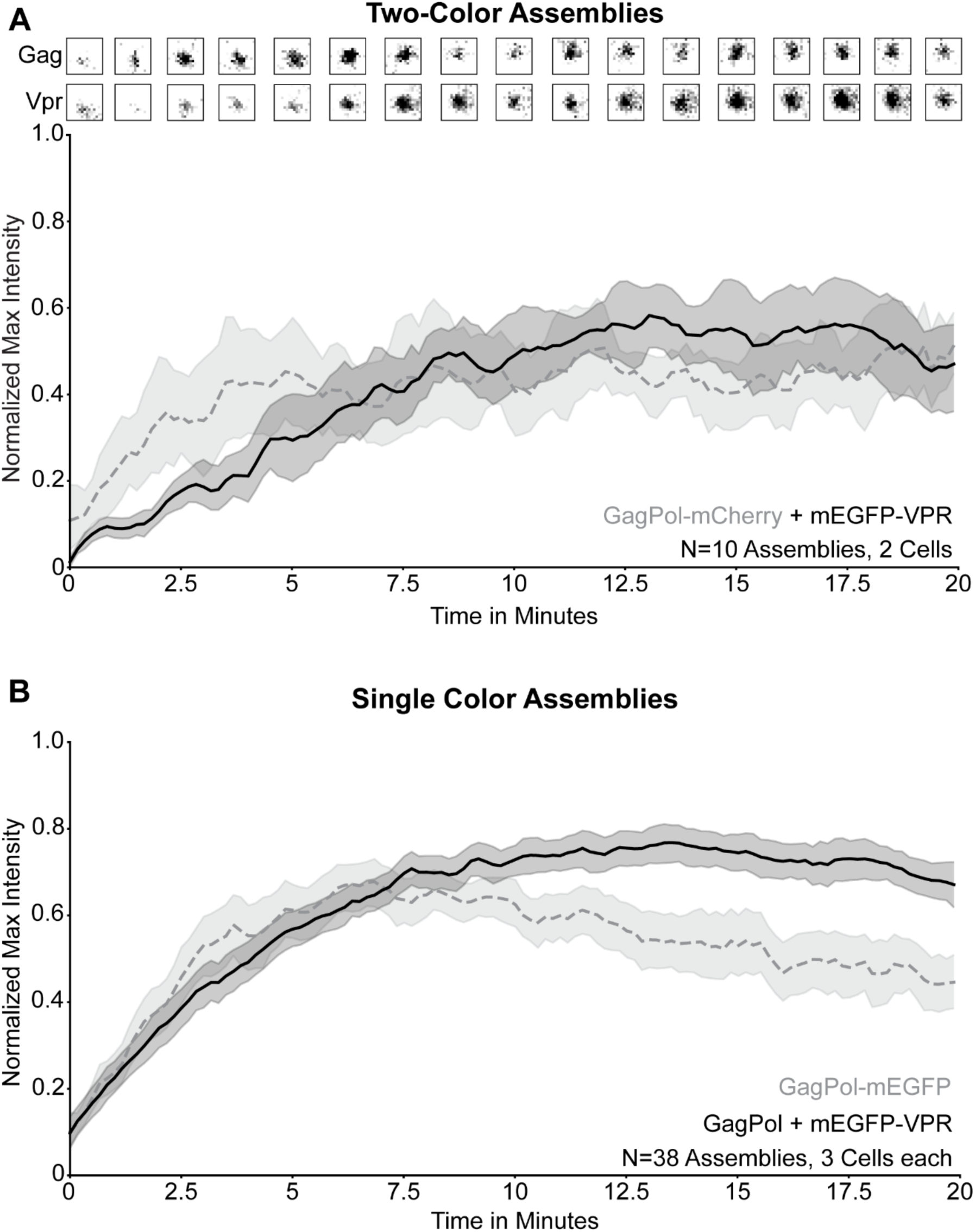
Assembly traces of GagPol and Vpr. **A** Assembly of Vpr and Gag-Pol. Top: select images from assembly. Bottom: The fluorescence intensity of the recruitment of mEGFP-Vpr and Gag-Pol-mCherry (average of 10 assemblies). **B** Traces for assembly traces for 38 VLPs where Vpr and GagPol were imaged separately in different cells. GagPol-mEGFP was imaged without Vpr. mEGFP-Vpr was imaged with untagged GagPol. The solid line in the graphs in both **A** and **B** shows the average while the shaded region surrounding the solid line shows the 95% confidence interval. Each assembly was rescaled to be on a 0-1 axis via (I-I_min_)/(I_max_-I_min_).

## Discussion

Our data shows that Vpr co-assembles with Gag, but with a slight delay. We also confirm that the LXXLF motif is required for Vpr recruitment. The delay of assembly of Vpr compared to Gag suggests that Vpr might be recruited to Gag only after Gag is at the membrane rather than Vpr being attached to Gag before arriving at to the membrane. It has been shown previously that Vpr and Tsg101 can competitively bind to Gag since their recruitment sites are close together (41). The competition with Tsg101 may also contribute to the delay of Vpr assembly compared to Gag. Our imaging can detect overall fluorescence, but it cannot determine whether there is turnover at the LXXLF binding site. It is possible that Tsg101 and Vpr might influence each other when it comes to binding Gag. This potential interaction of Tsg101/Vpr might also suggest why only ∼340 Vpr proteins are found in VLPs when approximately 2400 Vpr binding sites exist in a single assembled virus (4,19).

Our findings also suggest that Vpr is a good way to monitor assembly in-trans without tagging or altering Gag. Any protein fused to Vpr would arrive early in the assembly process, stay through scission, and be present in the final VLP. This contrasts with proteins like the ESCRTs which only show up in the final stages of assembly and don’t make it into the final VLP (5–8).

The incorporation kinetics of other accessory proteins like Vif and Nef are also unknown. It has been reported that Vif gets into particles by interactions with the viral RNA (42), and it is suspected that Nef gets into particles due to its myristoylation and interaction with membranes (16,43–46). Imaging could reveal other aspects of the kinetics of that packaging and give a fuller picture of how the HIV-1 virion is put together.

## Methods

### Cell Culture

HeLa and Hek293T cells were grown in Dulbecco’s Modified Eagle’s medium (DMEM, Gibco) supplemented with l-glutamine, sodium pyruvate, and 10% fetal bovine serum (FBS, Sigma). For live cell imaging HeLa cells were seeded onto MatTek dishes with no. 1.5 coverslips coated with fibronectin (Invitrogen). HeLa cells were transfected 6 hours before imaging with Fugene 6 (Promega) and 1000ng of DNA. Before imaging, media was replaced with cell imaging media (HBSS (Sigma), 10mM HEPES, pH7.4) supplemented with 1% FBS.

Hek293T cells were transfected with Lipofectamine 2000 (Thermo Fisher) and 1000ng of DNA 24 hours before VLP collection for imaging and 48 hours before VLP collection for westerns.

### VLP Collection and Imaging

Media from transfected Hek293T cells was collected either 24 or 48 hours after transfection and filtered through a 0.22µm filter (Millex). 1mL of cell media was carefully pipetted over a 20% sucrose in PBS solution and centrifuged at >30,000g for 1 hour. All supernatant was removed and collected VLPs were either resuspended in 100uL PBS for imaging or in 65ul of RIPA Buffer for westerns.

The collected VLPs in PBS were placed onto Poly-D-Lysine coated MatTek dishes with a no. 1.5 coverslip. VLPs were allowed to adhere for 20 minutes in the dark, then 2.5mL of PBS was added to each dish for imaging.

### Plasmids

GagPol-mCherry, GagPol-mEGFP, GagPol-LXXLF, and GagPol-mCherry-LXXLF were all created from the PCRV1-NL4.3-GagPol packaging vector which was a gift from the Paul Bieniasz at Rockefeller University. Both mCherry and mEGFP were cloned with no linker after the Matrix domain in Gag via Gibson assembly with the HiFi DNA Assembly cloning kit (New England Biolabs). Similar constructs have been previously published. The LXXLF mutation inserts a premature stop codon at the 43 amino acid within p6 which removes the last 10 amino acids of p6. This stop codon affects the Gag coding sequence, but makes no amino acid changes to the Pol reading frame. This mutation was created with the Quikchange site-directed mutagenesis kit (Agilent).

The mEGFP-Vpr construct was altered from the Addgene 110200 plasmid containing TEV-Vpr which was a gift from Sergi Padilla Parra (39). TEV was removed from the original plasmid and mEGFP was inserted in its place using a Gibson reaction via the HiFi DNA Assembly cloning kit (New England Biolabs).

### Biochemistry

Hek293T cell lysates were collected by adding 200uL RIPA buffer directly to cells after removing media. Cells were kept on ice for 20 minutes before RIPA was collected, vortexed, and frozen overnight. Collected VLPs and cell lysate were run on a 4-12% Tris-Glycine gel (Novex). Either an Anti-HIV-1 p24 Monoclonal (183-H12-5C) obtained from the NIH AIDS Reagent Program (47,48), a monoclonal anti-GFP (Living Colors JL-8, Clontech), or a monoclonal anti-β-actin (Sigma-Aldrich) primary antibody was used. HRP-coupled secondaries were used. Westerns were visualized on a LiCOR using ECL Prime.

### Assembly Selection and Plotting

Videos of assembly of Gag or Vpr in HeLa cells were analyzed with Metamorph software. The camera has 100 units added to each pixel and this we subtracted from all frames. From there, individual assemblies were selected and trimmed to the first time the assembling puncta was visible and the last time the puncta was visible. This automatically sets the first frame of the assembly as time zero. To account for the differing intensities between mEGFP and mCherry and for the differing intensities between different cells, the intensities for each assembly trace were rescaled by the following equation: 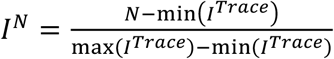. After rescaling, a rolling filter was applied to the traces. The rescaled max intensity for each frame of the assemblies were plotted together using python.

### VLP Quantification

VLPs from cells transfected only with GagPol-mCherry or GagPol-mCherry-LXXLF were collected and imaged with excitation at 488nm and 594nm (n=100). Puncta present when imaged with 594nm excitation were selected via with a 16×16 pixel circular region and the max intensities were recorded. The same regions were used to measure the max intensities during excitation at 488nm. The percentage of mCherry excitation at 488nm was calculated to be 20.551% of the excitation at 594nm.

For VLPs from cells transfected with GagPol-mCherry and mEGFP-Vpr or GagPol-mCherry-LXXLF and mEGFP-Vpr, puncta that were present when excited at 594nm were selected with circular regions and the max intensity was recorded. The same regions were then applied to an image taken with excitation at 488nm and the max intensities were recorded. The max intensities from excitation at 488nm had 20.551% of the max intensities with excitation at 594nm subtracted from them. If the resulting value of max intensity with 488nm excitation after subtraction was greater than zero, then that represented a red puncta that was also positive for green fluorescence.

### Statistics

P values for VLP quantification were calculated via a Chi-square test.

## Acknowledgements

We would like to thank Daniel Scott Johnson at Hofstra University and Joan Pulupa at Rockefeller University for help building and maintaining our microscope. We would also like to acknowledge the National Institute of General Medical Sciences of the National Institutes of Health for an award to SMS: R01GM119585.

